# Construction of a germline-specific RNAi tool in *C. elegans*

**DOI:** 10.1101/446898

**Authors:** Lina Zou, Di Wu, Xiao Zang, Zi Wang, Zixing Wu, Di Chen

## Abstract

Analysis of complex biological functions usually requires tissue-specific genetic manipulations in multicellular organisms. The *C. elegans* germline plays regulatory roles not only in reproduction, but also in metabolism, stress response and aging. Previous studies have used mutants of *rrf-1*, which encodes an RNA-directed RNA polymerase, as a germline-specific RNAi tool. However, the *rrf-1* mutants showed RNAi efficiency in somatic tissues. Here we constructed a germline-specific RNAi strain by combining an indel mutation of *rde-1*, which encodes an Argonaute protein that functions cell autonomously to ensure RNAi efficiency, and a single-copy *rde-1* transgene driven by the *sun-1* germline-specific promoter. The germline RNAi efficiency and specificity are confirmed by RNAi phenocopy of known mutations, knockdown of GFP reporter expression, as well as quantitative RT-PCR measurement of tissue-specific mRNAs upon RNAi knockdown. The germline-specific RNAi strain shows no obvious deficiencies in reproduction, lipid accumulation, thermo-tolerance and life span compared to wild-type animals. By screening an RNAi sub-library of phosphatase genes, we identified novel regulators of thermo-tolerance. Together, we have created a useful tool that can facilitate the genetic analysis of germline-specific functions in *C. elegans*.

## Introduction

The nematode *C. elegans* serves as a great model organism in biology research largely due to the ease of genetic manipulations. Genetic screens either by chemical mutagens or RNAi (double-stranded RNA-mediated gene silencing) have led to many discoveries. In *C. elegans*, effective RNAi knockdown can be achieved by feeding animals with *E. coli* that produce double-stranded (ds) RNAs corresponding to worm genes^1^. Genome-wide RNAi screens have been performed by many labs since the construction of the whole genome RNAi libraries, which are collections of *E. coli* strains that produce dsRNAs against nearly every gene in the *C. elegans* genome^2^. More focused RNAi screens are also applicable using RNAi sub-libraries of genes that encode transcription factors, chromatin-related factors, kinases, phosphatases and so on.

Numerous studies have demonstrated that multicellular organisms actively use across tissue communications to coordinate biological functions. Thus, tissue-specific genetic manipulations are frequently required to address complex biological questions. Researchers using *C. elegans* as a model have developed tools to perform tissue-specific RNAi experiments^3,4^. The strategies usually involve tissue-specific promoters-driving transgene rescue of mutations that are essential for the RNAi machinery. *rde-1*, which encodes an Argonaute protein, functions cell autonomously to ensure RNAi efficiency^5^. Therefore, tissue-specific promoters-driving *rde-1* rescue strains will allow RNAi to be effective in a tissue-specific manner.

The *C. elegans* germline plays regulatory roles in many biological processes. The germline not only serves as the reproductive tissue that produces gametes, but also affects metabolism, stress response and life span through non-autonomous regulation of gene expression in distal tissues^6-11^. However, the germline tissue is difficult for genetic manipulations since transgenes created by traditional methods are usually silenced in the germline. It was originally reported that mutations in RRF-1, an RNA-directed RNA polymerase, allow RNAi to be effective only in the germline but not in somatic tissues^12^. However, later studies revealed that the *rrf-1* mutants maintain RNAi efficiency in the soma, including tissues like the intestine and epidermis^13^.

In order to facilitate the genetic analysis in *C. elegans* germline, we set out to create a tissue-specific RNAi strain that allows RNAi to be functional effectively and specifically in the germline. Through CRISPR/Cas9-based genome editing and Mos1 transposon-based transgenic approaches, we constructed an indel mutation of *rde-1* that carries a single-copy *rde-1* transgene driven by the *sun-1* germline-specific promoter. The germline RNAi efficiency and specificity were validated by (1) RNAi phenocopy of known mutations, (2) knockdown of tissue-specific GFP reporter expression via *gfp* RNAi, and (3) quantitative RT-PCR measurement of tissue-specific mRNAs upon corresponding RNAi treatments. Furthermore, the germline-specific RNAi strain shows indistinguishable phenotypes in reproduction, neutral lipid accumulation, thermo-tolerance and life span when compared to wild-type animals. Lastly, we performed an RNAi sub-library screening of phosphatase genes in the germline-specific RNAi strain and identified novel regulators of thermo-tolerance. Together, we have created a useful tool that will help to analyze gene functions in *C. elegans* germline.

## Results

### Construction of a germline-specific RNAi strain via single-copy transgenic rescue of an *rde-1* indel mutation in the germline

In order to study gene functions in *C. elegans* germline, we sought to construct a germline-specific RNAi tool by transgenic rescue of an RNAi machinery mutant (Fig. 1A). Previous studies have applied similar approaches to construct epidermis, muscles and intestine-specific RNAi strains^3,4^. However, the *rde-1* deficiencies were either an E411K missense mutation^14^ or a Q825Ochre nonsense mutation^4^ that is close to the C terminus of the protein. These point mutations may not completely abrogate RDE-1 functions, which could lead to leakiness of RNAi efficiency in other tissues. To solve this problem, we used CRISPR/Cas9-based genome editing tools to create an *rde-1* indel mutation, which carries a 67-bp insertion and a 4-bp deletion in the exon 2 of the *rde-1* coding region. Although there is no frame shift in this allele, three premature stop codons (ochre) were introduced to the second exon, which is close to the beginning part of the coding sequence (Fig. 1B). Thus, this indel mutant is a potential null allele of *rde-1*.

**Figure 1.**
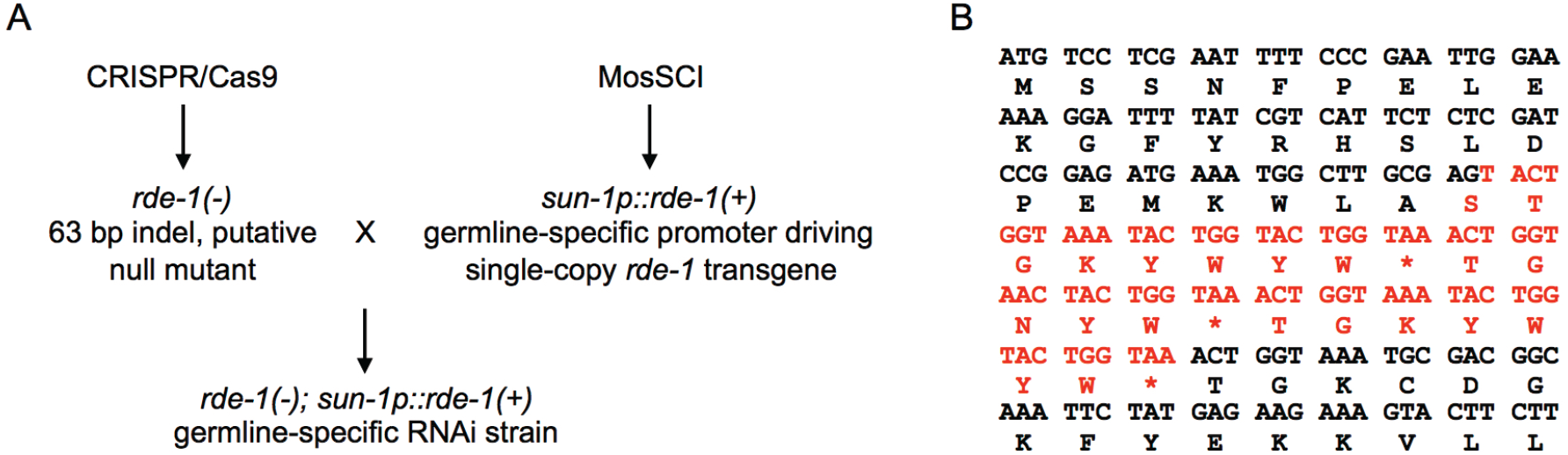
Experimental design of the germline-specific RNAi strain construction. (A) Workflow of the experimental design. An *rde-1* indel mutation was created using CRISPR/Cas9, and a single-copy *rde-1* transgene driven by the *sun-1* promoter was constructed via the MosSCI method. The *rde-1* mutation and germline *rde-1* transgene were then combined by genetic crosses. (B) cDNA and encoded amino acid sequences of the *rde-1* indel mutation. *, stop codon.

High copy number-transgenes produced by conventional methods, either in the form of extrachromosomal arrays or integrated, are prone to silencing in *C. elegans* germline. Therefore, we used the Mos1-mediated single copy insertion (MosSCI) method^15^ to make a single-copy *rde-1* transgene driven by the germline-specific *sun-1* promoter, which has been shown to be specifically and broadly active in the germline^16,17^. The *rde-1* mutant and transgene were then combined by genetic crosses to create the putative germline-specific RNAi strain.

### The germline-specific RNAi strain shows robust RNAi efficiency in the germline

To determine whether the *sun-1* promoter driving *rde-1* expression in the germline rescues the RNAi deficiency caused by the *rde-1* mutation, we treated the wild-type N2 and germline-specific RNAi strain with *gld-1* RNAi. *gld-1* encodes an RNA-binding protein that functions downstream of the GLP-1/Notch pathway to regulate the mitotic vs. meiotic fates of germline nuclei. Loss-of-function mutations in *gld-1* cause germline nuclei over-proliferation that leads to tumorous germline without oocytes^6,18,19^. Similar to wild-type animals, the germline-specific RNAi strain showed germline defects upon the *gld-1* RNAi treatment in about 50% penetrance (Fig. 2A, Table 1). We next tested the germline RNAi efficiency by knockdown of germline GFP expression via *gfp* RNAi. DEPS-1 is a P-granule-associated protein that is localized in the germline^20,21^. DEPS-1::GFP expression is significantly diminished by the *gfp* RNAi treatment in both the wild-type and germline-specific RNAi backgrounds (Fig. 2B). Finally, we performed RT-qPCR experiments to test the RNAi efficiency for genes expressed only in the germline, such as *pgl-1*^22^ and *daz-1*^23,24^. Knockdown of *pgl-1* or *daz-1* by RNAi effectively decreased mRNA levels of these genes in the germline-specific RNAi strain (Fig. 2C, D). Together, these results demonstrated the robust germline RNAi efficiency in the strain that we constructed.

**Figure 2.**
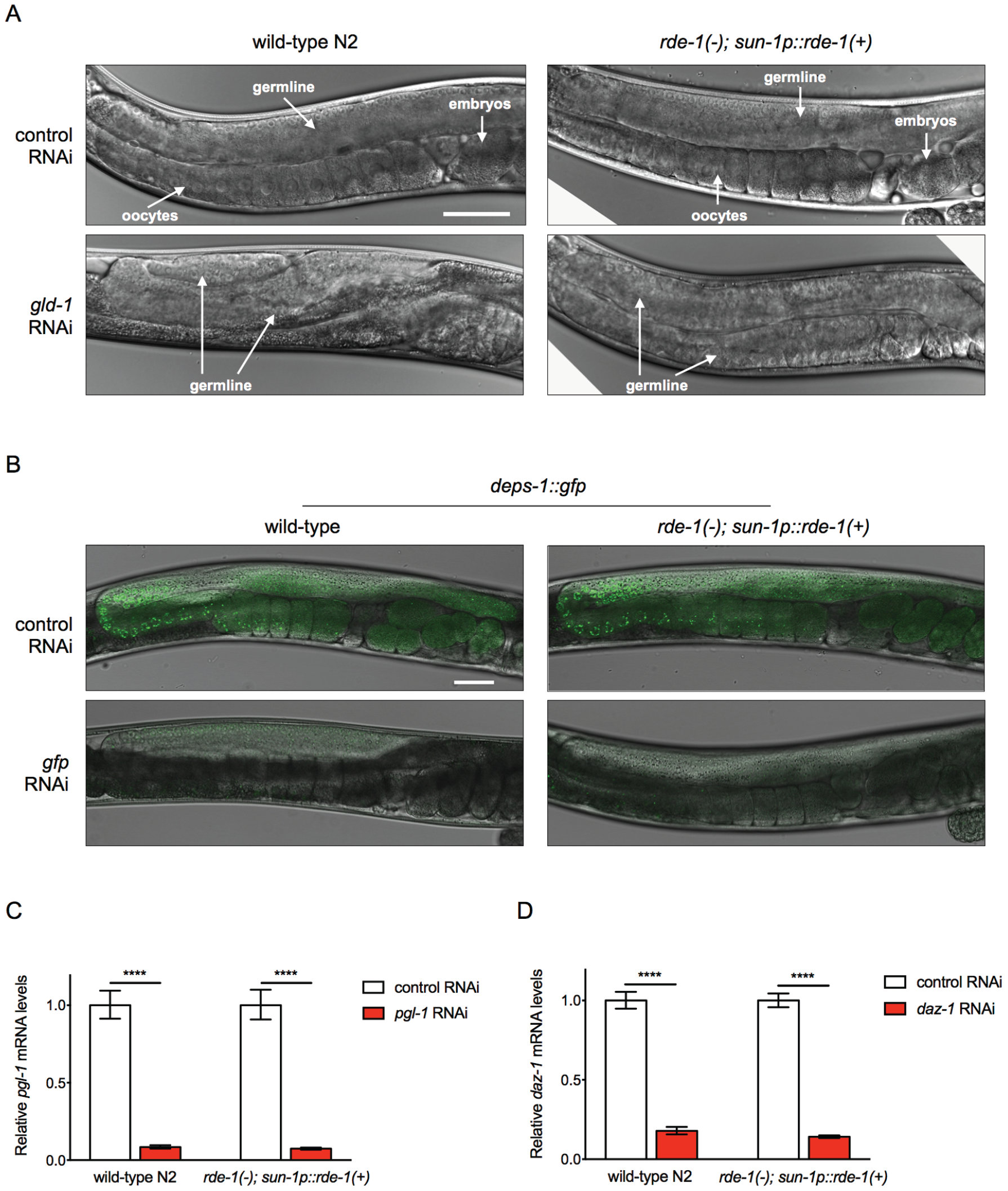
The germline-specific RNAi strain shows efficient RNAi knockdown in the germline. (A) Knockdown of *gld-1* leads to over-proliferating, tumorous germline phenotypes in the wild-type N2 and germline-specific RNAi strain. Scale bar, 50 μm. (B) DEPS-1::GFP expression in the germline can be effectively suppressed by *gfp* RNAi treatment in wild-type and germline-specific RNAi strain. Scale bar, 50 μm. (C-D) RT-qPCR measurement of germline genes *pgl-1* (C) and *daz-1* (D) mRNA levels showed effective knockdown by corresponding RNAi treatments in the wild-type N2 and germline-specific RNAi strain. ****, *p* < 0.0001 (*t* - tests).

**Table 1.** RNAi efficiency in different tissues in the germline-specific RNAi strain

| Tissues investigated | Genes targeted by RNAi | RNAi phenotypes | Numbers of animals <sup>a</sup> |  |
| --- | --- | --- | --- | --- |
|  |  |  | Wild-type | *rde-1(-); sun-1p::rde-1(+)* |
| Germline | *gld-1* | Pro, Tum | 42 / 98 | 51 / 102 |
| Germline | *egg-5* | Emb | 83 / 98 | 85 / 96 |
| Intestine | *elt-2* | Gob, Clr | 94 / 105 | 2 / 122 |
| Epidermis | *dpy-10* | Dpy | 130 / 140 | 6 / 114 |
| Epidermis | *tsp-15* | Bli, Dpy | 82 / 90 | 0 / 96 |
| Muscles | *unc-112* | Prz | 88 / 90 | 0 / 112 |
| Muscles | *pat-4* | Prz | 93 / 97 | 0 / 128 |

<sup>a</sup>Numbers of animals showing RNAi phenotypes vs. total numbers of animals scored.
Pro: proximal germ cell proliferation abnormal; Tum: tumorous germline
Emb: embryonic death
Gob: gut obstructed; Clr: clear
Dpy: dumpy
Bli: blistered
Prz: paralyzed

### The germline-specific RNAi strain shows no obvious RNAi efficiency in somatic tissues

The *rrf-1* mutant, which was initially used as a germline-specific RNAi tool^12^, shows leakiness of RNAi effects in the intestine and epidermis^13^. We then tested whether our germline-specific RNAi strain has RNAi efficiency in somatic tissues, including the intestine, epidermis and muscles. We first examined tissue-specific genes, RNAi knockdown of which show morphological phenotypes. Knockdown of ELT-2, an intestinal GATA transcription factor, leads to intestine developmental defects and a clear appearance^13,25^. *dpy-10* encodes a cuticle collagen, RNAi of which results in a Dpy (dumpy, short and fat) phenotype^26^. UNC-112 is a muscle dense body / M-line component. Knockdown of *unc-112* results in paralysis^27^. Unlike wild-type animals, the germline-specific RNAi strain did not show any of these phenotypes upon corresponding RNAi treatments (Fig. 3A, B). We next tested *gfp* RNAi efficiency using tissue-specific promoter driving GFP reporters. Compared to the wild-type N2, the *gfp* RNAi treatments could not reduce GFP expression produced by the intestinal *ges-1p::gfp*^28^ (Fig. 3C), epidermal *nlp-29p::gfp*^29^ (Fig. 3D) or muscular *myo-3p::gfp*^30^ (Fig. 3E) reporters in the germline-specific RNAi strain.

**Figure 3.**
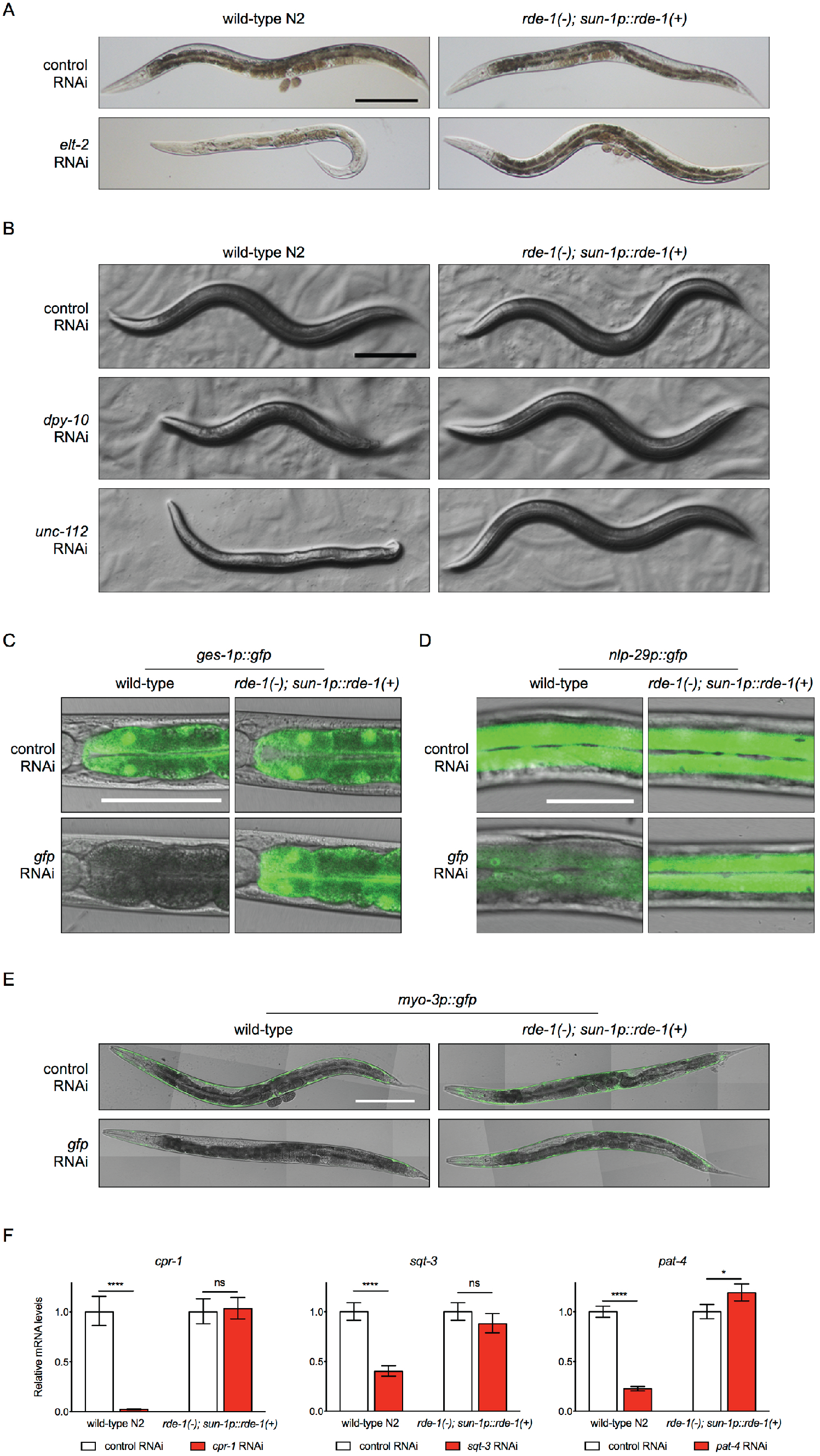
The germline-specific RNAi strain shows no obvious RNAi efficiency in somatic tissues. (A-B) The germline-specific RNAi strain showed no obvious RNAi phenotypes in the intestine by *elt-2* RNAi (A), in the epidermis by *dpy-10* RNAi and in the muscles by *unc-112* RNAi (B) compared to the wild-type N2. Scale bars, 200 μm. (C-E) The germline-specific RNAi strain showed no changes in tissue-specific promoters-driving GFP expression in the intestine (C), epidermis (D) or muscles (E) upon *gfp* RNAi treatments. (F) mRNA levels of tissue-specific genes *cpr-1* (intestine), *sqt-3* (epidermis) and *pat-4* (muscles) showed no reduction upon corresponding RNAi treatments in the germline-specific RNAi strain. ****, *p* < 0.0001; *, *p* < 0.05; ns, *p* > 0.05 (*t* - tests).

To quantitatively access the RNAi efficiency, we performed RT-qPCR experiments to measure mRNA levels of tissue-specific genes *cpr-1* (intestine)^31^, *sqt-3* (epidermis)^32^ and *pat-4* (muscles)^33^, upon corresponding RNAi treatments. Unlike wild-type animals, the germline-specific RNAi strain showed no obvious reduction of the tested mRNAs (Fig. 3F). Taken together, these data demonstrate that the germline-specific RNAi strain does not allow RNAi to be effective in the soma.

### The germline-specific RNAi strain shows no obvious deficiencies in reproduction, lipid metabolism, thermo-tolerance and life span

Since the germline-specific RNAi strain could be used to study regulatory roles of the germline on development, metabolism and aging, we examined whether this strain shows normal physiological features. The germline-specific RNAi strain has normal reproduction profile, total brood size and reproductive span compared to the wild-type N2 (Fig. 4A, B). Oil Red O staining with fixed animals and quantification indicate that the germline-specific RNAi strain has normal neutral lipids levels (Fig. 4C, D). Survival rate of animals treated with heat shock (35°C for 10 hours) and life span phenotypes are also indistinguishable from wild-type animals (Fig. 4E, F). Therefore, the germline-specific RNAi strain can be used to study the effects of germline knockdown in a variety of assays.

**Figure 4.**
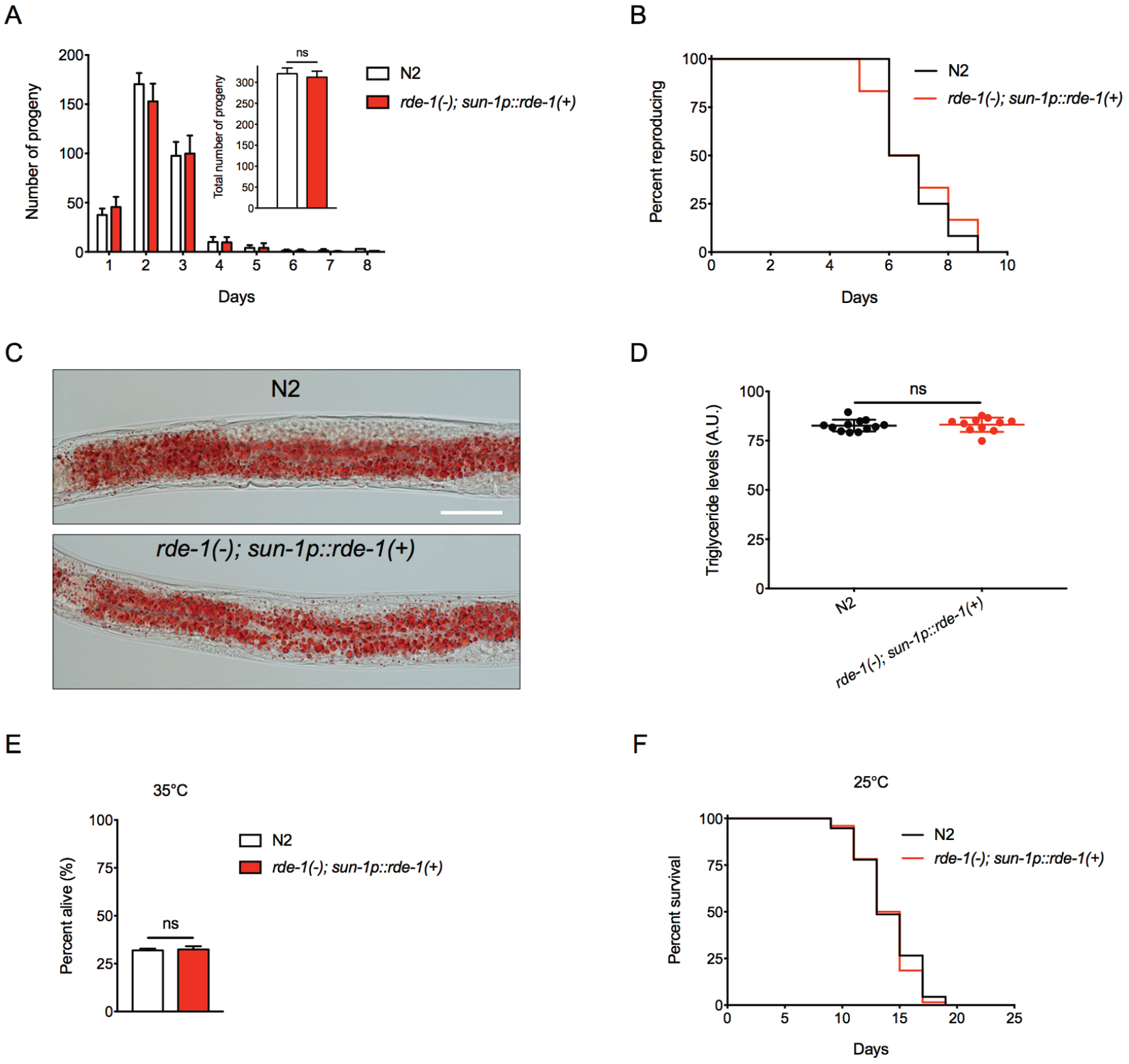
The germline-specific RNAi strain has normal reproduction, neutral lipids accumulation, thermo-tolerance and life span. (A) Reproductive profile and total brood size of the wild-type N2 and germline-specific RNAi strain. ns, *p* = 0.1513 (*t* - test). (B) Reproductive span of the wild-type N2 and germline-specific RNAi strain. *p* = 0.8527 (log-rank test). (C-D) Oil Red O staining (C) and quantification (D) of neutral lipids in the wild-type N2 and germline-specific RNAi strain. Scale bar, 50 μm. ns, *p* = 0.1858 (*t* - test). (E) Thermo-tolerance of the wild-type N2 and germline-specific RNAi strain at 35°C. ns, *p* = 0.6986 (*t* - test). (F) Lifespan of the wild-type N2 and germline-specific RNAi strain at 25°C. *p* = 0.3543 (log-rank test).

### Identification of novel regulators of thermo-tolerance by an RNAi-based genetic screen

One important purpose of constructing the germline-specific RNAi strain is to use this tool for RNAi-based genetic screens. As a proof of principle experiment, an RNAi screen was performed to identify novel regulators of thermo-tolerance, since increased intrinsic thermo-tolerance has been associated with the delay of aging^34^. Phosphatases in many cases play regulatory roles in various signal transduction pathways. However, most of the *C. elegans* phosphatase genes have not been well characterized for their biological functions, especially in a tissue-specific context. Thus, we chose an RNAi sub-library containing RNAi against 163 phosphatase genes to perform the screen for increased thermo-tolerance in the germline-specific RNAi background. After the primary screen and several rounds of re-tests, we identified four phosphatase genes *R155.3*, *paa-1*, *W01B6.6* and *upp-1*, RNAi knockdown of which leads to significantly improved thermo-tolerance (>10%, *p* < 0.001) compared to the control RNAi treated animals (Fig. 5A). R155.3 and W01B6.6 are predicted tyrosine phosphatases. PAA-1 is the sole *C. elegans* homolog of PR65, a subunit of the protein phosphatase 2A (PP2A) complex^35^. UPP-1 is a uridine phosphorylase, mutations in which result in resistance to the anticancer drug 5-fluorouracil^36^. None of these phosphatases have been associated with heat stress response, especially in a tissue-specific manner.

**Figure 5.**
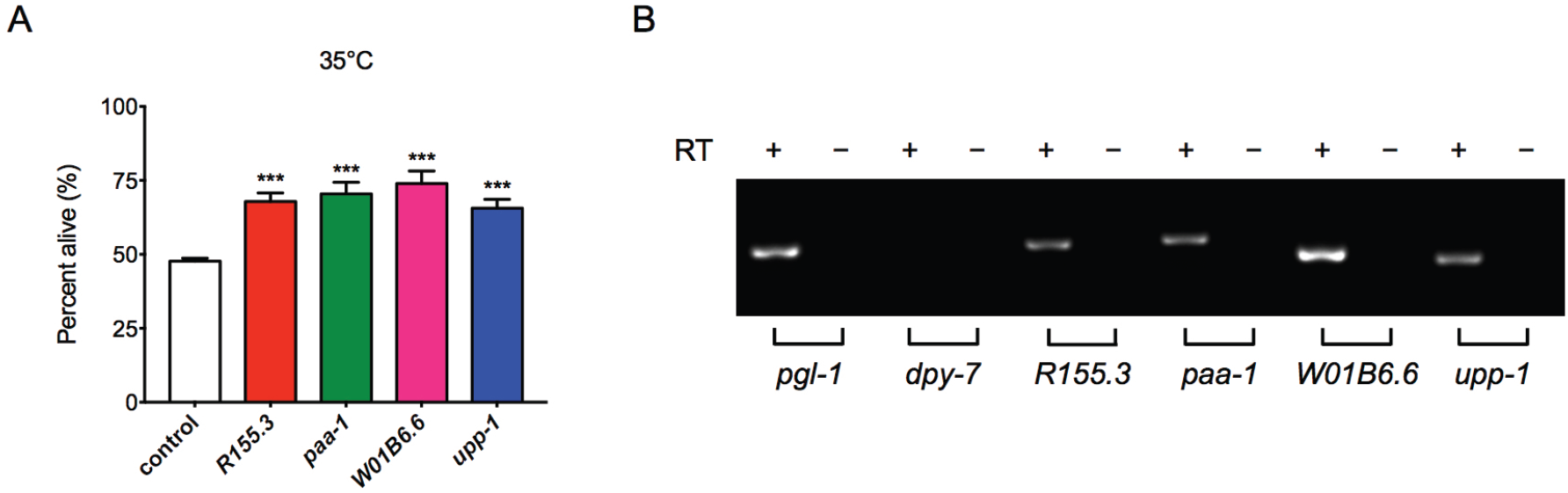
Identification of novel regulators of thermo-tolerance by a germline-specific RNAi screen of phosphatase genes. (A) Germline knockdown of phosphatases encoding genes *R155.3*, *paa-1*, *W01B6.6* and *upp-1* significantly increases survival at 35°C. ***, *p* < 0.001 (*t* - test). (B) RT-PCR products of *R155.3*, *paa-1*, *W01B6.6* and *upp-1* amplified from reverse transcription products using RNAs extracted from dissected gonadal tissues. *pgl-1* (germline gene) and *dpy-7* (epidermal gene) serve as positive and negative controls, respectively. RT, reverse transcriptase.

In order to examine whether the identified phosphatase genes are expressed in the germline, we performed RT-PCR experiments to detect transcripts of these genes with RNAs extracted from micro-dissected gonadal tissues (Fig. 5B). *pgl-1*, a germline gene^22^, and *dpy-7*, an epidermal gene^37^, were used as the positive and negative controls, respectively. Transcripts of *R155.3*, *paa-1*, *W01B6.6* and *upp-1* can be detected by RT-PCR (Fig. 5B), suggesting they are expressed in the germline.

## Discussion

The ease of gene knockdown by RNAi and the existence of whole genome RNAi libraries have enable researchers to perform exploratory studies using *C. elegans* as a model. The tissue-specific RNAi strains, which allow RNAi to be effective only in certain tissues, are very useful tools to dissect genes’ functions. The *C. elegans* germline has many biological functions. Besides the critical role in reproduction, it can also affect metabolism, stress response and aging at the whole animal level. In many cases, the regulatory mechanisms involve across tissues, endocrine-like signaling. However, the existing germline-specific RNAi strains all show RNAi efficiency in the soma. Considering the importance of germline in various biological processes, we set out to make a tool that allows RNAi to be effective specifically in the germline.

The Argonaut protein RDE-1, a key component of the RNA-induced silencing complex (RISC), is required for RNAi to be effective^5^. Since RDE-1 functions cell autonomously, tissue-specific transgenic rescue of the *rde-1* mutation will make RNAi to be effective in a spatially restricted manner^4,14^. Thus, we made a germline-specific *sun-1* promoter driving *rde-1* single copy transgene to rescue the *rde-1* loss-of-function mutation. Initially, we used the *rde-1(ne219)* mutant, which carries a E411K missense mutation, for the strain construction. This mutation has been used to construct epidermis and muscle-specific RNAi strains in previous studies^14^. Further characterization of this strain (DCL484 *mkcSi13 [sun-1p::rde-1::sun-1 3’UTR + unc-119(+)] II; rde-1(ne219) V*) revealed that it shows RNAi leakiness in the soma (Fig. S1). We speculated that the leakiness was caused by the residue activities of the *rde-1(ne219)* mutation. Thus, we used CRISPR/Cas9 to create an indel *rde-1* mutation and re-constructed the germline-specific RNAi strain (DCL569 *mkcSi13 [sun-1p::rde-1::sun-1 3’UTR + unc-119(+)] II; rde-1(mkc36) V*). Further analyses of this strain demonstrated that it does not show obvious RNAi activities in the soma in various assays (Fig. 3). Therefore, it might worth re-constructing the existing tissue-specific RNAi strains using the null allele of *rde-1* instead of the *rde-1(ne219)* hypomorphic allele to achieve better tissue-specific knockdown.

Besides RNAi knockdown, researchers in the community have created other tools to achieve tissue-specific genetic manipulations. For example, researchers have adapted the auxin inducible degradation (AID) system in *C. elegans*^38^. AID allows drug-inducible, tissue-specific depletion of proteins. However, it requires knock-in of a Degron encoding sequence to the genes of interests via CRISPR/Cas9, which makes this technique more appropriate for focused studies rather than large scale genetic screens. The germline-specific RNAi strain created in this study is useful for high throughput genetic studies. A pilot RNAi screen of phosphatase genes for thermo-tolerance phenotypes were performed, and several novel heat stress resistance regulators were identified. These results demonstrate that the germline-specific RNAi tool that we constructed can be used for genetic screens to identify biological functions of germline genes.

## Methods

### *C. elegans* strains and maintenance

Strains were cultured on NGM agar plates seeded with *E. coli* OP50 at 20°C unless otherwise stated. The following *C. elegans* strains were obtained from the Caenorhabditis Genome Center:

Bristol (N2) strain as the wild-type strain

CB7272 *ccIs4251 [(pSAK2) myo-3p::GFP::LacZ::NLS + (pSAK4) myo-3p::mitochondrial GFP + dpy-20(+)] I; mIs12 [myo-2p::GFP + pes-10p::GFP + F22B7.9p::GFP] II; dpy-17(e164) III; frIs7 [nlp-29p::GFP + col-12p::DsRed] IV; uIs69 [pCFJ90(myo-2p::mCherry) + unc-119p::sid-1] V*

EG4322 *ttTi5605 II; unc-119(ed9) III*

JH3207 *deps-1(ax2063[deps-1::GFP]) I*

MAH23 *rrf-1(pk1417) I*

SJ4144 *zcIs18 [ges-1::GFP(cyt)]*.

The following strains were generated in D.C. lab:

DCL455 *mkcSi13 [sun-1p::rde-1::sun-1 3’UTR+ unc-119(+)] II; unc-119(ed3) III*

DCL484 *mkcSi13 [sun-1p::rde-1::sun-1 3’UTR + unc-119(+)] II; rde-1(ne219) V*

DCL565 *rde-1(mkc36) V*

DCL569 *mkcSi13 [sun-1p::rde-1::sun-1 3’UTR + unc-119(+)] II; rde-1(mkc36) V*

DCL588 *ccIs4251 [(pSAK2) myo-3p::GFP::LacZ::NLS + (pSAK4) myo-3p::mitochondrial GFP + dpy-20(+)] I*

DCL589 *frIs7 [nlp-29p::GFP + col-12p::DsRed] IV*

DCL590 *ccIs4251 [(pSAK2) myo-3p::GFP::LacZ::NLS + (pSAK4) myo-3p::mitochondrial GFP + dpy-20(+)] I; mkcSi13 [sun-1p::rde-1::sun-1 3’UTR] II; rde-1(mkc36) V*

DCL582 *mkcSi13 [sun-1p::rde-1::sun-1 3’UTR + unc-119(+)] II; frIs7 [nlp-29p::GFP + col-12p::DsRed] IV; rde-1(mkc36) V*

DCL592 *deps-1(ax2063[deps-1::GFP]) I; mkcSi13 [sun-1p::rde-1::sun-1 3′UTR + unc-119(+)] II; rde-1(mkc36) V*

DCL593 *mkcSi13 [sun-1p::rde-1::sun-1 3′UTR + unc-119(+)] II; rde-1(mkc36) V; zcIs18 [ges-1::GFP(cyt)]*.

### Molecular cloning

The *sun-1* promoter driving *rde-1* rescue plasmid was constructed by cloning PCR fragments of the 468 bp *sun-1* promoter, 3572 bp *rde-1* genomic sequence with all the exons and introns, and 780 bp *sun-1* 3′UTR into the MosSCI vector pCFJ151 (Addgene #71720) using the NEBuilder HiFi DNA Assembly Cloning Kit (NEB).

The *rde-1* konckout plasmid was generated by inserting targeted sgRNA fragments into the pDD162 vector (Addgene #47549) using the site-directed mutagenesis kit (TOYOBO). The sgRNAs were designed using the CRISPR DESIGN tool (http://crispr.mit.edu).

sgRNA 1 sequence: TTATCGTCATTCTCTCGATC

sgRNA 2 sequence: AGGCCCACTGGTAAATGCGA

### Generation of the *rde-1* indel mutation by CRISPR/Cas9

The *rde-1(mkc36)* indel mutation was generated via CRISPAR/Cas9-based genome editing approach^39^. A DNA mix containing the Cas9-sgRNA plasmids (50 ng / μl) and selection marker pCFJ90 P*myo-2*::mCherry (2.5 ng / μl) was injected into N2 young adults. Animals from the F1 generation were screened by PCR for insertions and / or deletions. The *rde-1(mkc36)* homozygous mutant was identified from the F2 generation by PCR and the mutations were verified by DNA sequencing of PCR products.

### Single-copy transgene by MosSCI

The *sun-1p::rde-1* transgenic strain was constructed by injection of a DNA mix containing 37.5 ng / μl targeting plasmids, 50 ng / μl pCFJ601 (P*eft-3*::Mos1 Transposase), 10 ng / μl neuronal selection marker pGH8 (P*rab-3*::mCherry) and 2.5 ng / μl pharyngeal selection marker pCFJ90 (P*myo-2*::mCherry) into the EG4322 strain (*ttTi5605 II; unc-119(ed9) III*) according to the protocol previously described^15^. After injection, worms were maintained at 25°C until starved. Single non-Unc worms without the selection markers were spread to new plates. Successful insertions were confirmed by PCR and DNA sequencing.

### RNAi by feeding

*E. coli* strains that carries either the empty vector L4440 (control) or various gene-targeting constructs were cultured and induced for dsRNA production as previously described^40^. For RNAi treatments, gravid adult worms were allowed to lay eggs on RNAi plates at 20°C for 2 hours. Synchronized progenies were collected for various assays at day 1 of adulthood. All RNAi clones were verified by DNA sequencing.

### Reproduction profile

L4 larvae were individually placed onto NGM plates and then transferred to new plate every 24 hours at 20°C. Numbers of progeny on each plate were counted 2 days later after removing the adult animals.

### Lipid staining by Oil Red O

Oil red O staining was performed as previously described^41^. L4 larvae were collected and fixed in 1% formaldehyde and frozen at −80°C. The samples were frozen in dry ice / ethanol bath and thawed under a stream of warm water for three cycles. After washing twice with the S buffer, worms were incubated with Oil red O (3 mg / ml) solution for 30 min at room temperature. Animals were then washed with the S buffer and incubated on ice for 15 min. Images were taken with a Nikon Eclipse Ni-U microscope and DS-Fi2 color CCD. Triglyceride levels of the second pair of intestinal cells were quantified using the Image J software.

### Thermo-tolerance

Synchronized day 1 adult worms were incubated at 35°C for 10 hours before counting the numbers of alive or dead animals. About 80-100 worms were used in each experiment and each assay was repeated for three times.

### Life span

Worms at the late L4 stages transferred to fresh NGM plates and incubated at 25°C for survival. FUdR (20 μg / ml) was added onto NGM plates during the reproduction period to prevent progeny production. Animals were scored as alive, dead (no response to gentle touch) or lost (death from non-aging causes) every other day. Survival curves were plotted with Prism 6 software and statistical analyses were performed using the log-rank method.

### RNAi screen

An RNAi sub-library that contains 163 predicted phosphatase genes was used to identify regulators of thermo-tolerance. Synchronized DCL569 *mkcSi13 [sun1p::rde-1::sun-1 3′UTR + unc-119(+)] II; rde-1(mkc36) V* L1 larvae were transferred onto the seeded RNAi plates and incubated at 20°C until animals reached the day 1 adulthood. In the primary screen, RNAi-treated animals were incubated at 35°C for 12 hours until the control RNAi treated animals were all dead. Plates with live animals were regarded as thermo-tolerant candidates and the corresponding genes were re-tested in the secondary screen. Candidates that passed two rounds of tests were used in the final assays, in which RNAi-treated animals were incubated at 35°C for 10 hours, and the survival percentages were compared with the control RNAi-treated animals.

### RT-qPCR

About 600 synchronized day 1 adult worms were collected and frozen in the Trizol reagents (Takara). Total RNAs were extracted using the Direct-zol RNA mini prep kit (ZYMO Research) and the cDNAs were synthesized using the reverse transcription system (Takara). Quantitative PCR reactions were performed in triplicates on a Roche LightCycler 480 real-time PCR machine using the SYBR Green dye (Takara). Relative gene expression levels were calculated using the 2^−ΔΔCt^ method^42^. RT-qPCR experiments were performed at least three times with independent RNA extractions.

### RT-PCR using dissected gonadal tissues

L4 larvae were transferred into the S buffer (100 mM NaCl and 50 mM potassium phosphate [pH 6.0]) on a glass slide. Heads of animals were cut off near the pharynx using syringe needles to collect the gonad. About 40-50 gonadal tissues were collected in the Trizol reagents for total RNA extraction.

## Acknowledgements

We would to thank members of the Di Chen lab for discussions, and Dr. Zhuo Du for strains, Drs. Mengqiu Dong and Wenhong Zhang for advice on CRISPR/Cas9 genome editing experiments. Some strains were provided by the CGC, which is funded by NIH Office of Research Infrastructure Programs (P40 OD010440). D.C. is supported by the National Natural Science Foundation of China (31471379, 31671527) and Natural Science Foundation of Jiangsu, China (BK20141316).

## Author contributions

Conceived and designed the experiments: L.Z. and D.C. Performed the experiments: L.Z., D.W., X.Z., Z.W. and Z.W. Analyzed the data: L.Z. and D.C. Wrote the paper: L.Z. and D.C. All authors reviewed the manuscript.

---

**Figure S1.**
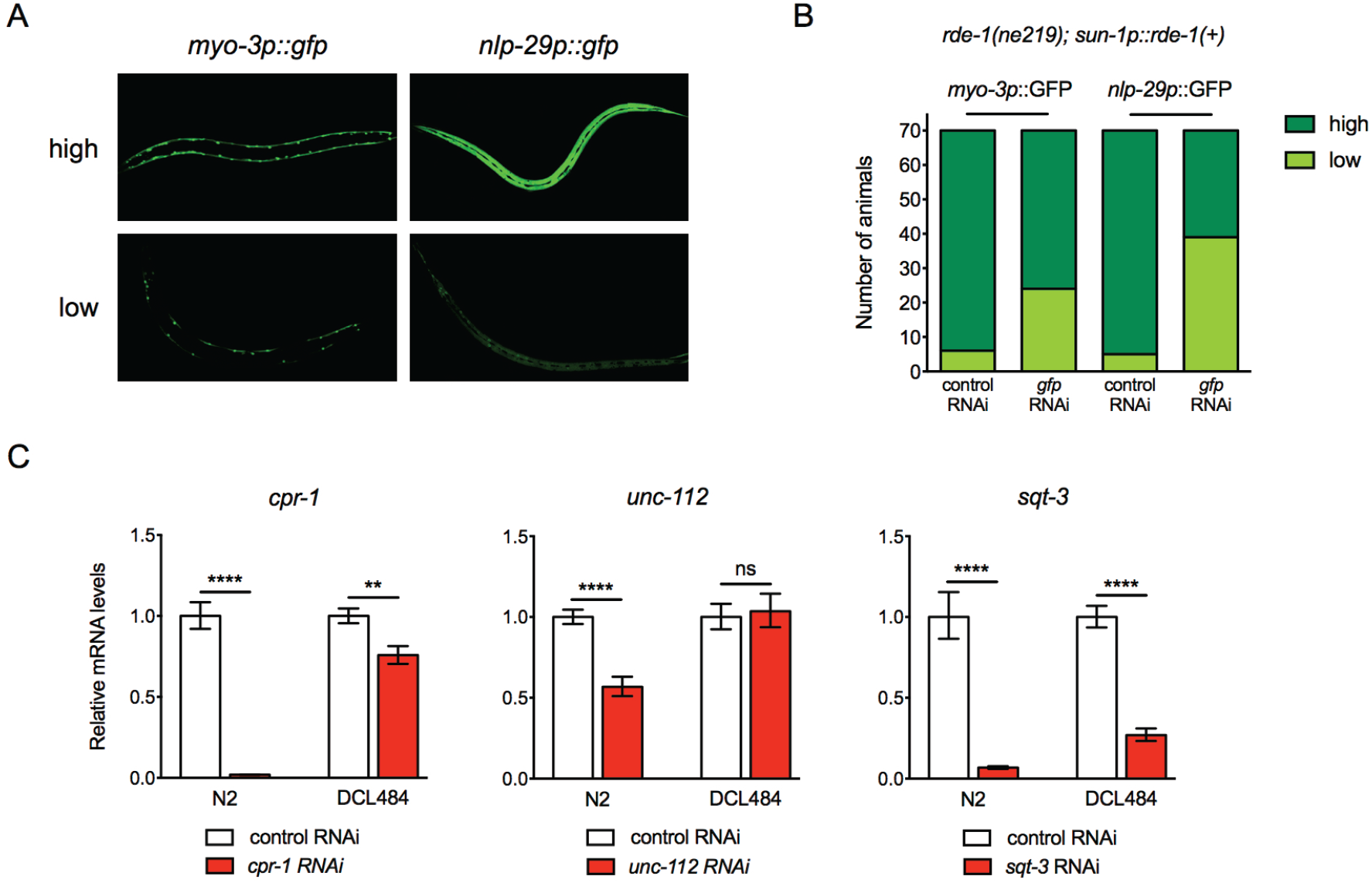
The DCL484 germline RNAi strain shows RNAi efficiency in the soma. (A) Representative images of muscular (*myo-3*) and epidermal (*nlp-29*) GFP reporters treated with *gfp* RNAi in the DCL484 germline RNAi strain, which carries the *rde-1(ne219)* hypomorphic mutation. (B) Quantification of GFP expression. 70 animals were scored in each treatment. (C) mRNA levels of tissue-specific genes *cpr-1* (intestine) and *sqt-3* (epidermis) showed significant reduction upon corresponding RNAi treatments in the DCL484 germline RNAi strain. ****, *p* < 0.0001; **, *p* < 0.01; ns, *p* > 0.05 (*t* - tests).

**Table S1.**
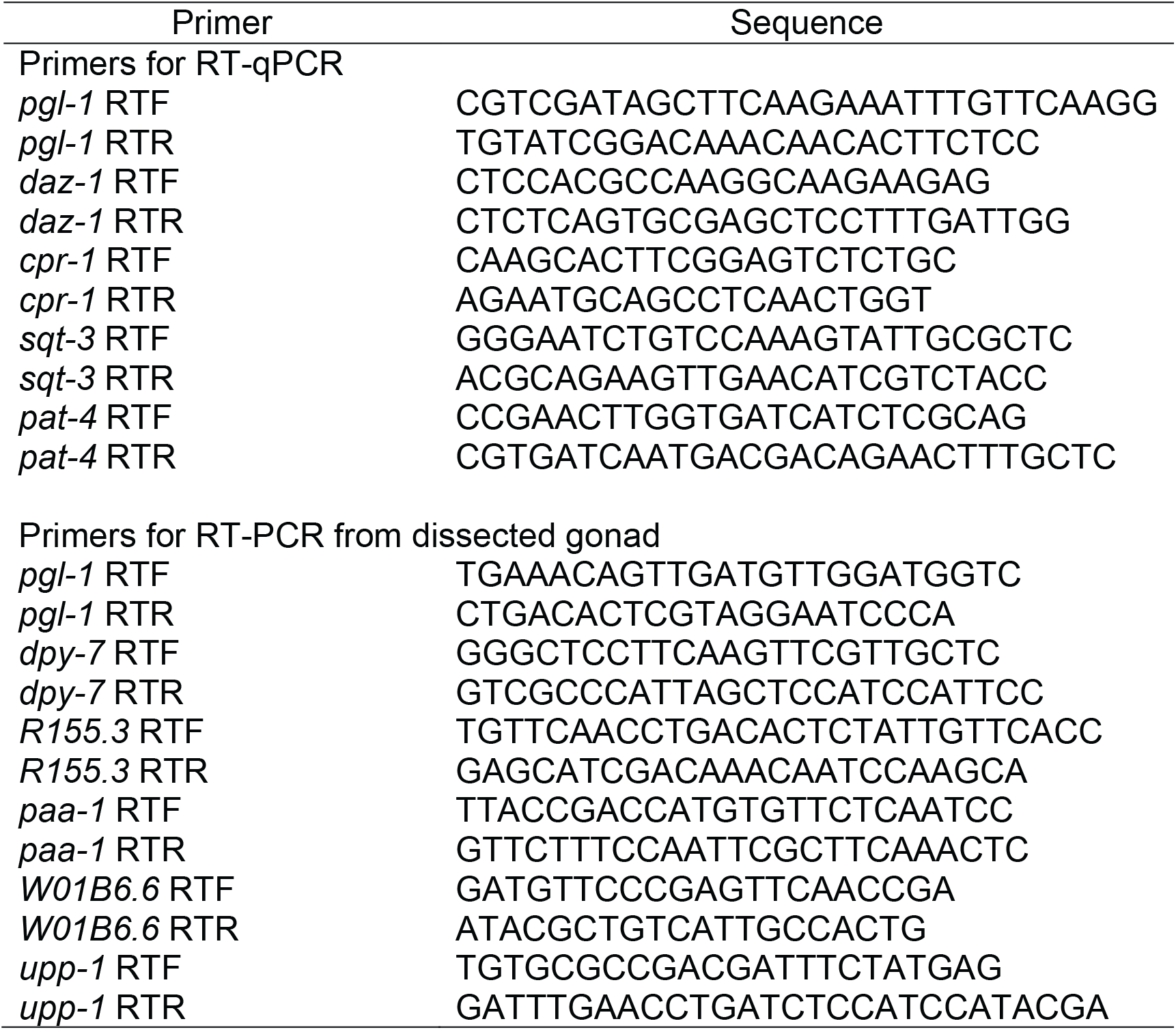
Sequences of RT-PCR primers

